# Predicting the Risk of Avian Influenza Zoonosis using Viral Genome Sequencing Data

**DOI:** 10.64898/2026.08.21.746166

**Authors:** Anna Fairweather, Austin Andrews, Joshua Grier, Liam Brierley, Lorenzo Cattarino, Jasmina Panovska-Griffiths

## Abstract

Avian Influenza viruses (AIVs) infect a broad host range despite having a natural reservoir in wild aquatic birds. Whilst most strains stay within their host species, some break the species barrier through genetic adaptations. We are most concerned about zoonotic cases, where a human becomes infected. Despite these events being rare, they are associated with high mortality and introduce the risk of onward human-to-human transmission of AIV. As a novel pathogen within the human population, this could have pandemic potential.

Using genetic composition features for 8 AIV proteins drawn from viral sequence data, we employ machine-learning algorithms to classify AIV cases as zoonotic or not. These genetic features encode host ‘signatures’ which can indicate zoonosis and include frequency measures such as dipeptide composition and amino acid physiochemical properties.

We consistently find XGBoost to outperform all other algorithms. We optimise parameters for ten classification models: one for each of the 8 proteins and two combined models. Following this, we show that a multi-model approach gives the best performing prediction for AIV zoonosis. We have identified all 8 proteins as having a role in predicting zoonotic transmission. Of particular importance is the PB2 and HA proteins, with specific amino acid physiochemical properties such as charge, secondary structure and hydrophobicity amongst the most indicative features in our combined models.

Our alignment-free computational study can identify AIV cases still within avian hosts which are genetically closest to zoonotic AIV cases, thereby identifying the cases most likely to cross the species barrier. In a resource limited environment, our model could be used to quickly identify high priority cases for further investigation.

**Key points:**

- XGBoost is the best performing machine learning algorithm for the binary classification task of identifying avian influenza cases as zoonotic or not.
- Our model suggests that all eight proteins we considered have a role in determining whether avian influenza cases have zoonotic potential or not.
- Genetic composition features of avian influenza proteins can encode information that can be used as zoonotic signatures.
- We have created a system that identifies high priority cases still within avian hosts that should be prioritised for further investigation.

## 1. Introduction

Avian Influenza viruses (AIVs) are enveloped, single stranded negative-sense RNA viruses of the Orthomyxoviridae family that can cause severe lower respiratory tract infections. Wild aquatic birds are considered to be the natural reservoirs of avian influenza, although AIVs are able to infect a broad range of hosts including avian, human and swine. Whilst most strains stay within their host species, infrequent genetic adaptations in multiple loci of the virus genome can cause a strain to cross the species barrier [1]. This transmission can be zoonotic (i.e. an Avian Influenza case identified within a human host). Although these zoonotic events are relatively uncommon, they are associated with high mortality and hospitalisation rates in humans, making them a cause for concern [2]. There is also potential for onward human-to-human transmission of AIV following a zoonotic infection. This could have pandemic potential, given that the virus would spread as a novel pathogen against which the general population has little-to-no immunity [3].

Influenza virus’ can be categorised into 4 types: A, B, C and D. Type A is the most common type amongst wild avian hosts. It has the greatest genetic variation and broadest host range, making it the type of highest concern. The type A genome is segmented into 8 RNA segments which encode over 10 viral proteins. These can be categorised as surface glycoproteins (HA and NA), matrix proteins (M1 and M2), polymerase basic proteins (PB1 and PB2), nucleoproteins (NP), polymerase acidic proteins (PA) and non-structural proteins (NS1 and NS2). Antigenic differences in the hemagglutinin (HA) and neuraminidase (NA) proteins are used to categorise Influenza A Virus’ (IAVs), with the type of HA and NA providing the namesake of IAV subtypes. These proteins cover around 80% and 17% of the virus surface respectively [4]. As a result, they play a vital role in Influenza transmission.

The primary purpose of HA is to mediate fusion between the Influenza viral envelope and the host cell membrane via a receptor binding [5]. This gives it a key role in replication. The host tropism of IAV is thought to be primarily determined by its affinity to bind to specific sialic acid (SA) sites on host cells [6]. AIVs preferentially attach to SA via 2,3-SA linkage where as human IAVs prefer to bind to SA via *α*2,6-SA linkage [7].

It has been known for some time that swine strains of Influenza express preference for and contain both linkages [8]. Originally it was thought that humans and birds exclusively expressed *α*2,6-SA and *α*2,3-SA linkage respectively, and only swine expressed both. However, more recently, it has been found that many birds and mammals, including humans, have both types of receptors in varying quantities and in different tissues [9]. As a result, birds have now been identified as having a high probability of being “mixing vessels” for AIV and IAV in humans. Turkeys, chickens, quails and ducks have been identified as being a particular risk group for this due to their frequent human-to-bird contact [10].

Changes in the host tropism of AIV are determined by an accumulation of mutations in the viral genes which encode proteins, known as antigenic shift and drift [3, 11]. Due to the pandemic potential of AIV, there has been a focus on looking at genetic sequences of AIV to try and highlight strains of concern and develop a full understanding of the conditions needed for zoonotic transmission. This work has been primarily focussed on establishing specific amino acid (AA) substitutions and residues within different proteins that affect transmission. Supervised and unsupervised machine-learning algorithms (MLAs) can be developed and deployed to identify more of these AA substitutions.

The application of MLAs in medicine and biology is a growing field. Historically, these higher-order statistical models have been used to uncover information within neurological images [12], in cardiovascular risk assessment [13] or respiratory medicine [14]. More recently, MLAs have started to be used in population health and epidemiological research in research [15]. MLAs can also be used to identify genetic composition features that may encode host ‘signatures’, such as AA physiochemical properties. As part of this work, we have completed an extensive literature review on this topic; details can be found in the supplementary material Appendix A.1. We used this review to inform the methods we have used in our study.

The aim of our study is to address the gaps in application of MLAs to classify zoonotic avian influenza using genomic data. Our work will complement the range of avian influenza host tropism existing MLAs, but will also allow us to explore false positive prediction. This would provide a potential focus area for non-zoonotic cases in avian hosts which might be capable of making the zoonotic jump, and hence act as early warning of avian influenza cases which may soon become zoonotic. This could help to prevent sporadic zoonotic outbreak cases and, in the most extreme case, prevent an outbreak with pandemic potential.

We develop and apply a variety of MLAs to establish an optimal zoonosis prediction model, for each individual protein considered, as well as two combined models to compare their accuracy in classifying zoonosis. Using that model, we will then identify the key features that characterise avian influenza zoonotic transmission.

## 2. Methods

### 2.1. Data

The data consisted of 19,529 preprocessed avian influenza genome sequences collected from avian (18,911) and human (618) hosts. The raw sequences were sourced from GISAID and GenBank and belonged to 59 different avian influenza subtypes; details can be found in related work [16]. These raw sequences were processed into genome feature frequency measurements for 8 avian influenza proteins. In total, there were 7 different categories of genome feature frequency measurement.

These were: Nucleic Acid Composition of length 2, Dipeptide Composition (DPC), Composition-Transition-Distribution Composition (CTDC), Transition (CTDT), Distribution (CTDD), Conjoint Triad (CTriad) and Pseudo-Amino Acid Composition (PAAC). These feature groups measure different types of frequency measures of nucleic acids, amino acids and groups of amino acids with shared physiochemical properties. Detailed descriptions of these measurements can be found in Appendix A.2.

There are many features within each group, varying from 16 to over 400. In total, we have 8,440 genomic features for each sample of Avian Influenza, resulting in over 160 million data points in our data set.

We note that our data was quite sparse, with 46% of data points in our dataset entered as zero. We removed any features with more than 90% zeros and reran our models. This did not affect our results in any significant way; it did not improve any of our classification models. As a result, we retained all features despite some being sparse.

### 2.2. MLAs selection

There are numerous machine learning algorithms which are able to perform classification tasks that could be suited to our classification problem. Before we began optimising the parameters of any of our models, we did a preliminary test to see how different MLAs performed on the full dataset with their default parameters. We considered a number of MLAs that were selected following our scoping literature review (see supplementary material Appendix A.1): XGBoost, Random Forest, Logistic Regression, Ridge Classification, Decision Tree Classification, Support Vector Classification, k-Nearest Neighbours, Gaussian Naive Bayes, Neural Network and Quadratic Discriminant Analysis.

### 2.3. Training and testing MLAs

For each MLA, we normalised and one-hot encoded our data. Following this, we divided the genomic features dataset into separate training and testing dataset, by randomly selecting 20% of the samples to be the testing dataset. All training of prediction models was done with 5-fold cross-validation. This involves splitting the dataset into four training subsets and one testing subset, all of equal size. The training process involves iterating through a different subset as the testing subset, 5 times in total so that every sample is tested once and only once. This helps to prevent over fitting and combat the issue of an imbalanced data set.

### 2.4. Evaluating MLAs performance

We evaluated the performance of all our prediction models using a number of measures: accuracy, precision, F1 score, recall, specificity, Matthew’s correlation coefficient (MCC), area under the curve (AUC) and average precision (see Appendix A.3 for precise definitions). Due to our imbalanced data set, we believe that the most indicative and important performance measures are the F1 score, followed by recall. F1 score helps to account for the imbalanced data set and does not give an overinflated measure of performance due to the large number of avian cases (18,911) relative to the number of zoonotic cases (618). Furthermore, recall encourages false positivity above false negativity. It is more useful to have false positivity when trying to build a preventative predictive tool as this indicates cases which are not yet zoonotic but are likely to make the zoonotic jump in the near future. This is where policy makers most likely want to focus resources to try and contain these cases before they become zoonotic.

Comparing the accuracy across all MLAs, combined with the finding of our literature review, allowed us to select one optimal MLA best suited for the zoonosis classification task.

### 2.5. Parameter Optimisation

To achieve the best performance when training with the optimal MLA, we performed a parameter optimisation using a combination of grid search and random search.

Grid search involves testing all combinations of parameters in a predetermined parameter optimisation grid. It is a fully exhaustive algorithms that gives the best performing set of parameters once complete. Random search involves sampling random combinations of parameters within predetermined ranges. This is a more efficient method than grid search and allows higher dimensions of parameters to be searched. However, it is not exhaustive and could lead to the most efficient selection of parameters remaining unknown.

### 2.6. Feature prediction from the optimal MLA

Using the optimal MLA, we then trained 8 separate models using only the features that belonged to one of each of the 8 proteins in our dataset. This meant that we had 8 different predictive MLAs, each based on a different protein’s features. Following this, we took the top 20 significant features of each protein model (using SHAP) and used just these 160 features in another model which we call the ‘selected features combined model’. The aim of this was to see if we could reduce the complexity and running time of a model that used all the features without reducing performance significantly. Lastly, we ran a model using the entire dataset which we call the ‘all features combined model’.

For each of these variations of the optimal MLA, we identified the key genetic features that can indicate avian influenza zoonosis.

## 3. Results

### 3.1. Comparison of Classification Algorithms

A summary of the performance of each of the MLAs can be found in Table 1. Whilst a number of classifiers performed well, XGBoost outperforms all other classification algorithms in most metrics. Specifically, XGBoost achieved an F1 score of 0.8766 and a recall of 0.8188. Logistic Regression and Random Forest also performed notably well. However, unlike for XGBoost, we did not find a significant increase in their F1 score performance following parameter optimisation. Hence, we chose XGBoost as the optimal MLA, and proceeded to perform rigorous parameter optimisation on it and use it for feature characterisation.

**Table 1:** Summary of the initial performance when trained on the entire dataset of each tested classification algorithm.

| Algorithm | Accuracy | Precision | F1 Score | Recall | Specificity | MCC |
| --- | --- | --- | --- | --- | --- | --- |
| XGBoost | 0.9927 | 0.9450 | 0.8766 | 0.8188 | 0.9984 | 0.8757 |
| Random Forest | 0.9905 | 0.9362 | 0.8329 | 0.7509 | 0.9983 | 0.8336 |
| Logistic Regression | 0.9892 | 0.8178 | 0.8328 | 0.8496 | 0.9938 | 0.8277 |
| Ridge Classifier | 0.9816 | 0.7037 | 0.7135 | 0.7250 | 0.9900 | 0.7045 |
| Decision Tree Classifier | 0.9860 | 0.7762 | 0.7799 | 0.7849 | 0.9925 | 0.7730 |
| Support Vector Classifier | 0.9849 | 0.9015 | 0.7121 | 0.5891 | 0.9979 | 0.7218 |
| k-Nearest Neighbours | 0.9876 | 0.8891 | 0.7800 | 0.6959 | 0.9971 | 0.7802 |
| Gaussian Naive Bayes | 0.9620 | 0.4415 | 0.5343 | 0.6863 | 0.9710 | 0.5309 |
| Neural Network | 0.9864 | 0.8143 | 0.7282 | 0.7043 | 0.9956 | 0.7361 |
| Quadratic Discriminant Analysis | 0.9354 | 0.2862 | 0.4058 | 0.6975 | 0.9432 | 0.4204 |

### 3.2. Parameter Optimisation

We initially optimised over a large number of parameters for XGBoost. Following this initial optimisation with random search, we further fine-tuned the parameters using grid search. The parameters optimised with this more exhaustive method are the learning rate, maximum depth and number of estimators, since our initial large parameter scope identified them as the most important parameters to focus on. Table 2 shows the final optimised parameters for each individual protein model and the combined models.

**Table 2:** Optimised XGBoost parameters for all protein and combined models, using random search and grid search.

| Model | Learning Rate | Max Depth | Number of Estimators |
| --- | --- | --- | --- |
| HA | 0.55 | 9 | 110 |
| M1 | 0.25 | 7 | 120 |
| NA | 0.40 | 7 | 120 |
| NP | 0.25 | 7 | 120 |
| NS1 | 0.25 | 7 | 120 |
| PA | 0.25 | 7 | 120 |
| PB1 | 0.25 | 9 | 110 |
| PB2 | 0.25 | 9 | 110 |
| All Features Combined Model | 0.30 | 6 | 100 |
| Selected Features Combined Model | 0.30 | 6 | 100 |

### 3.3. Prediction Model Performance

Optimising parameters of the individual protein models led to good performance across the board for all proteins (see Table 3). The HA and PB2 protein models were particularly strong performers with F1 scores of 0.8103 and 0.8412, respectively. Despite this strong performance, no single protein model alone could outperform the combined models. The all features combined model outperformed all other models considered, with an F1 score of 0.8766. As it includes all features in the original dataset, it is the most computationally expensive. Figure 1 shows a confusion matrix of the all features combined model, showing its predictive capability over the 5 cross-validation folds.

**Table 3:** Performance of individual protein models and combined models using XGBoost with parameters optimised to each model.

| Model | Accuracy | Precision | F1 Score | Recall | Specificity | MCC | AUC | Average Precision |
| --- | --- | --- | --- | --- | --- | --- | --- | --- |
| HA | 0.9889 | 0.8810 | 0.8103 | 0.7525 | 0.9966 | 0.8080 | 0.9833 | 0.8582 |
| M1 | 0.9878 | 0.8765 | 0.7878 | 0.7186 | 0.9966 | 0.9809 | 0.9966 | 0.8113 |
| NA | 0.9880 | 0.8659 | 0.7956 | 0.7363 | 0.9962 | 0.7924 | 0.9812 | 0.8641 |
| NP | 0.9873 | 0.8797 | 0.7757 | 0.6959 | 0.9968 | 0.7756 | 0.9783 | 0.8296 |
| NS1 | 0.9882 | 0.8958 | 0.7921 | 0.7104 | 0.9973 | 0.7919 | 0.9795 | 0.8357 |
| PA | 0.9870 | 0.8752 | 0.7695 | 0.6878 | 0.9968 | 0.7692 | 0.9789 | 0.8362 |
| PB1 | 0.9872 | 0.8733 | 0.7756 | 0.7008 | 0.9966 | 0.7751 | 0.9807 | 0.8475 |
| PB2 | 0.9907 | 0.9183 | 0.8412 | 0.7767 | 0.9977 | 0.8398 | 0.9886 | 0.9018 |
| All Features Combined Model | 0.9927 | 0.9450 | 0.8766 | 0.8188 | 0.9984 | 0.8757 | 0.9925 | 0.9271 |
| Selected Features Combined Model | 0.9924 | 0.9419 | 0.8702 | 0.8091 | 0.9984 | 0.8691 | 0.9917 | 0.9246 |

**Figure 1:**
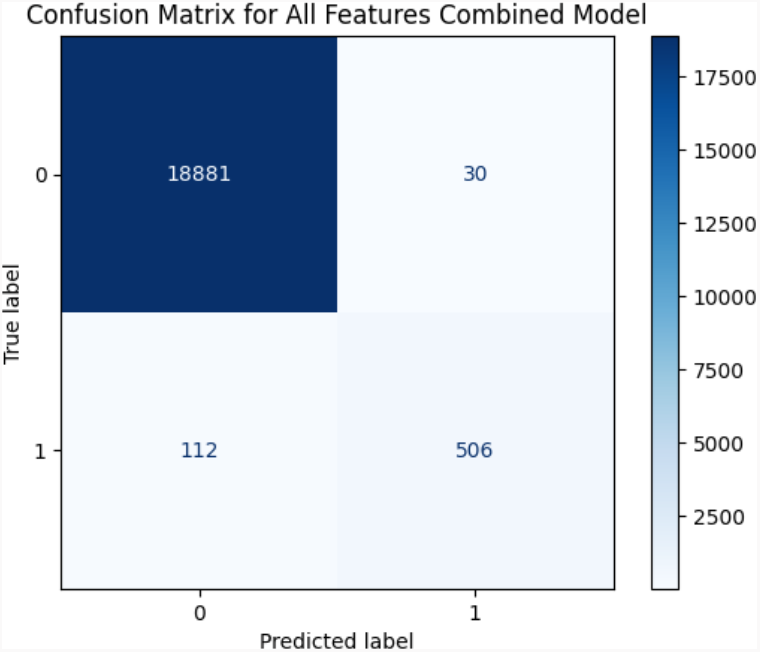
Confusion Matrix for the all features combined model over the 5 cross-validation folds. The indicators 0 and 1 refer to non-zoonotic and zoonotic, respectively.

Interestingly, the selected features combined model had only narrowly worse performance than the all features combined model (F1 score of 0.8702). This model takes only the top 20 features of each of the protein models (160 features total), resulting in it having significantly reduced computational complexity. Whilst this shows that having the full dataset does give the best performance, it also presents an opportunity to substantially reduce model complexity and run time without significantly affecting model performance.

In general, the nucleic acid composition features were one of the main contributing features across all of the individual protein models. A summary of the top 3 contributors for each model is given in Table 4. Notably, in both the all features combined model and the selected features combined model, the top contributing features involved far fewer nucleic acid composition features than the top contributing features for the individual protein models. Given the increased performance within these combined models, this suggests that the nucleic acid composition features are not actually more important than any other feature type when trying to determine if a case is zoonotic.

**Table 4:**
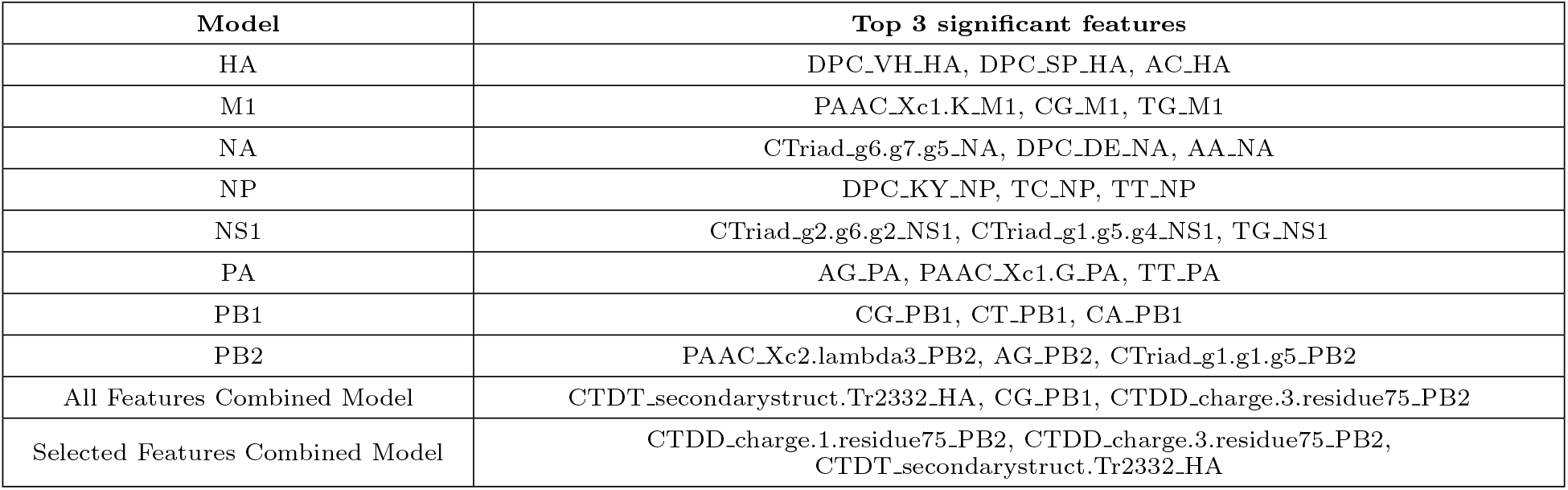
Top 3 contributing features for each of the individual protein models and the combined models.

Figure 2 shows bar charts of the top 30 contributing features to the all features combined model (a) and the selected features combined model (b). We discuss the physical relevance of a selection of these features more in Section 3.4. For now, we identify that there is overlap between these two feature lists. For example, the top 3 most significant features in the all features combined model are all within the top 7 features of the selected features combined model. While not all features appear in both lists, there is a lot of overlap which is reassuring. Furthermore, the differences in the top feature lists suggest that each model potentially offers new information that we are unable to derive from the other model, as both models are capturing slightly different features which are potentially related biologically to the host tropism of AIV cases.

**Figure 2:**
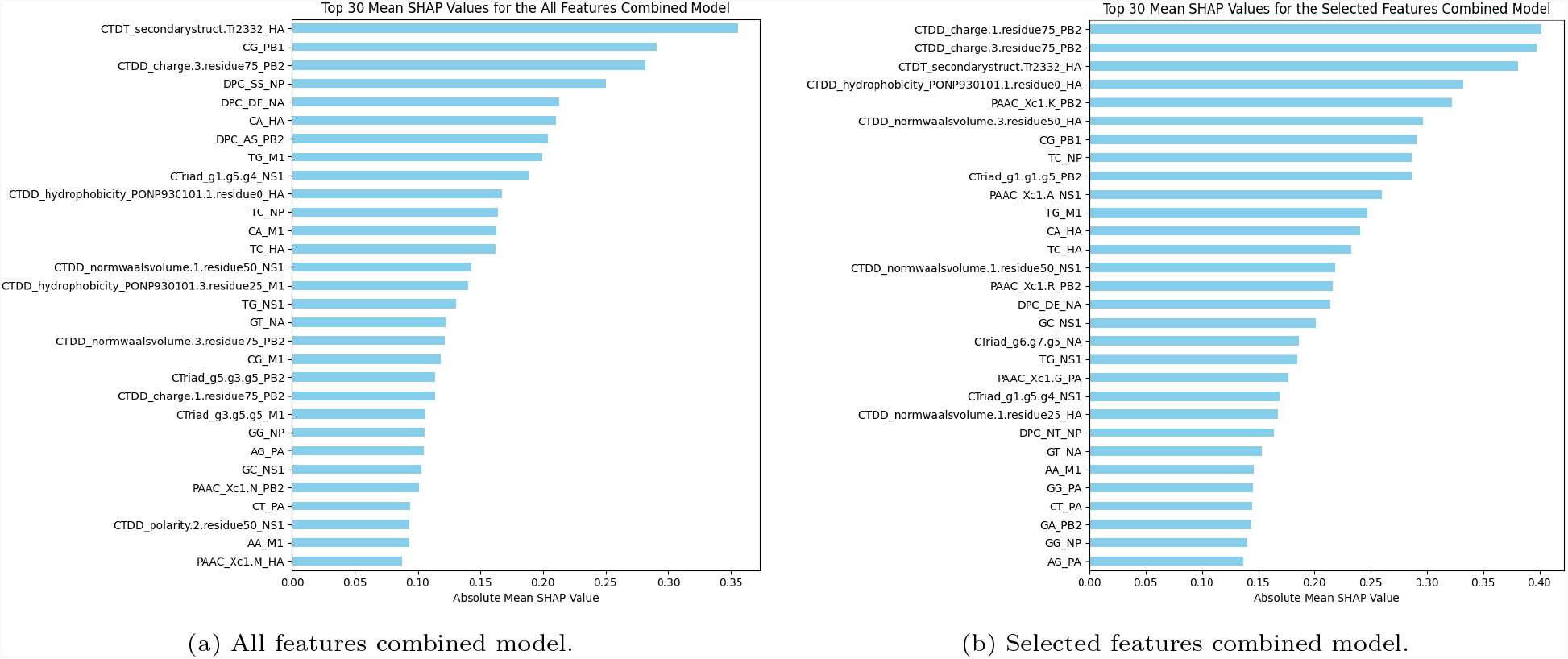
Top 30 mean absolute SHAP values for the (a) all features combined model and (b) selected features combined model.

One of the benefits of having individual protein models as well as combined models is that we can collate these together and compare how each case is being predicted across models. In Figure 3a we can see the binary prediction of all confirmed zoonotic cases across all the models. Despite many cases having zoonotic signatures across all models (indicated in red), some only have zoonotic signatures in just a few of the 10 models. There are even some zoonotic cases which have no zoonotic signatures in any models. Furthermore, there are some zoonotic cases that have no zoonotic signatures in any of the individual protein models whilst having a signature in one/both of the combined models. This shows that having zoonotic potential is likely not just given by the structural features of just one avian influenza protein. Instead, it likely involves many proteins and specific mechanisms of interaction between these proteins.

**Figure 3:**
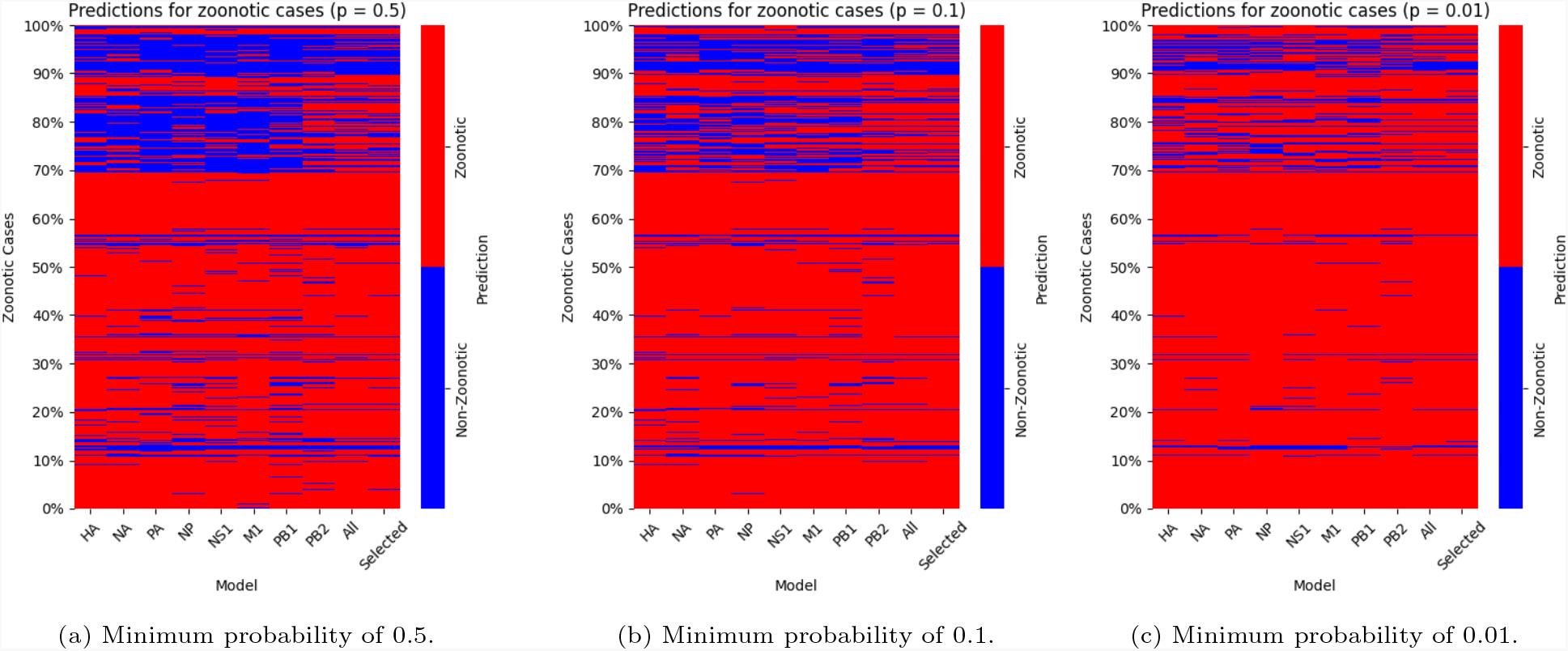
Prediction of zoonotic signatures of all confirmed zoonotic cases across all models using a minimum predicted probability of being zoonotic of (a) 0.5, (b) 0.1, and (c) 0.01.

We have further extended our model to include any prediction as zoonotic if its probability of being predicted as zoonotic is at least 0.1 and 0.01 (compared to a default probability 0.5 in the binary prediction). This can be seen in Figure 3b and Figure 3c respectively. We see that following this approach, there are fewer zoonotic cases that have no zoonotic signatures across models, improving the model’s predictive capability for the confirmed zoonotic cases. Especially when we set the probability as 0.01, we see over 98% of zoonotic cases have at least one zoonotic signature across all models. Moreover, setting the probability to be 0.1 or 0.01 does not have a large impact on the number of non-zoonotic cases being incorrectly predicted (compare Figure 4a to Figure 4b and Figure 4c). Even in the 0.01 case, less than 12% of non-zoonotic cases have at least 1 zoonotic marker. This is a useful result, as it suggests that our model could be used to identify non-zoonotic cases which are most at risk of becoming zoonotic, thereby offering a route of prioritisation of cases for further investigation. In conclusion, we should have some falsely predicted non-zoonotic cases as these likely indicate cases which are genetically closest to making the zoonotic jump.

**Figure 4:**
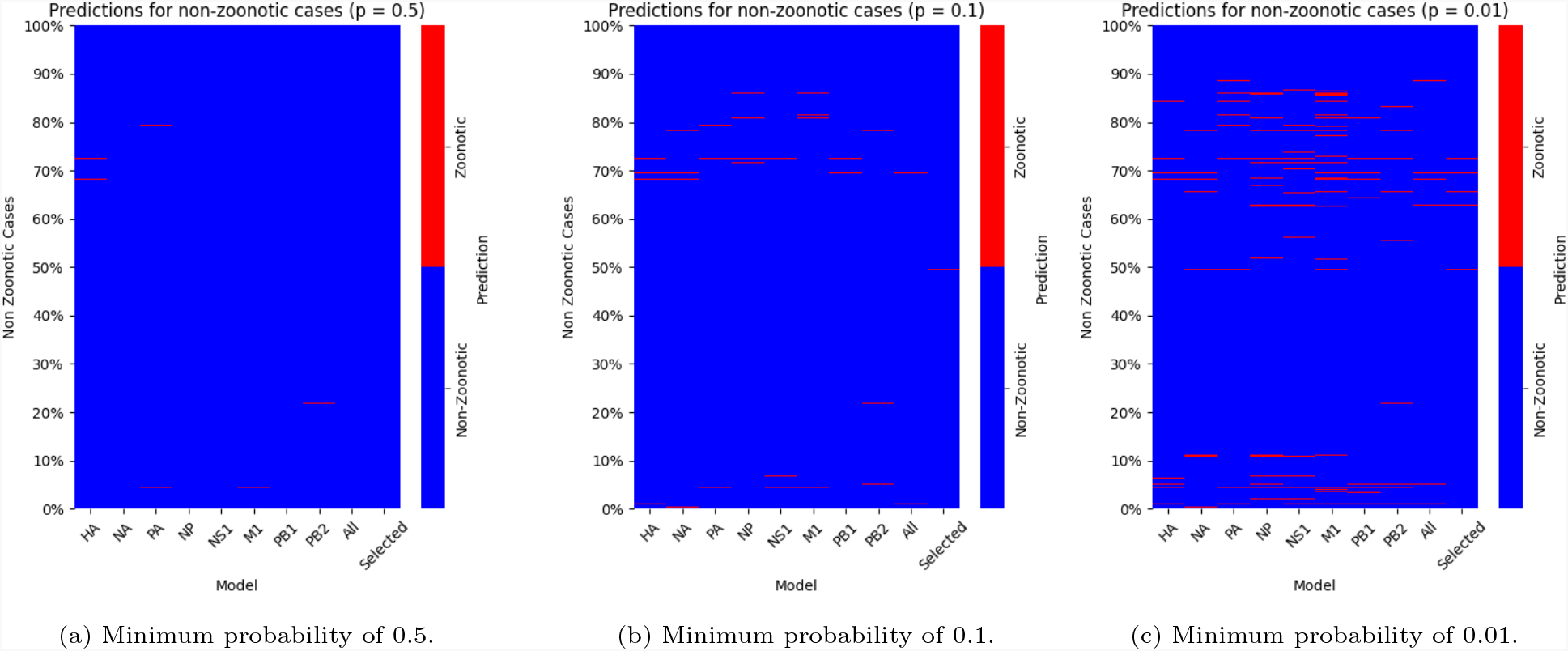
Prediction of zoonotic signatures of all non-zoonotic cases across all models using a minimum predicted probability of being zoonotic of (a) 0.5, (b) 0.1, and (c) 0.01.

### 3.4. Biological Interpretation

Whilst we do not give specific biological reasons for why our model has identified certain features as important, here we discuss the general biological interpretation of the roles of the proteins and some of their known biological features that influence transmission and, ultimately, zoonosis.

The HA protein provides the most important feature in the all features combined model, and 3 of the top 6 features in the selected features combined model. Furthermore, the individual HA protein model is one of the best performing individual protein models, falling only behind PB2 when comparing the F1 score. The HA protein mostly consists of a globular head and a stalk region. The globular head contains a receptor binding site (RBS) that comprises a small group of AAs. This site is surrounded by more AA structures whose length and composition vary with receptor binding preference. Specific AA substitutions in the RBS have been identified as determiners of receptor-binding specificity and have contributed to the emergence of previous AIV pandemics [11, 17, 18, 19]. In some subtypes, these substitutions have created a hydrophobic region which changes the glycan binding profiles. This hydrophobic region is incompatible with the hydrophilic *α*2,3-SA, but complementary to the hydrophobic *α*2,6-SA [3, 20].

Humans spread IAV through airborne routes whereas AIVs primarily spread through faecal routes between birds. Tosheva et al. [21] found that for IAV to be airborne transmitted, three phenotypic viral properties must change: receptor-binding specificity, polymerase activity and HA acidic stability. The first of these two have been well studied but the latter is less understood. We know that the membrane fusion process of HA is influenced by the host cell membrane pH and the acidic and thermal stability of HA. Stability here is defined as the ability to resist inactivation when exposed to extreme pH/temperatures. AA substitutions in HA have been identified as likely causing changes in its acidic stability [22, 18] as well as its thermal stability [19]. Human influenza viruses have a lower fusion pH than avian and swine viruses (pH 5.0-5.5 vs 5.6-6.0 and 5.5-5.9 respectively) [21]. These findings show that the acidic stability of the HA protein has an important role in influenza viral transmissibility, although changes in HA acidic stability are not enough alone for airborne transmissibility.

The best performing individual protein model, when measured by F1 score, was the PB2 model. The PB2 protein is responsible for some of the most significant features in both combined models. The polymerase proteins (PB1, PB2 and PA) are responsible for the transcription and replication processes of the viral RNA. AA mutations in PB2, especially E627K, have been identified in zoonotic transmission events [17, 23]. AA substitutions in polymerase proteins increase their activity and flexibility during transcription and replication. Multiple adaptive mutations have also been identified in PA and PB1, but the mechanisms undergone by these proteins are less clearly characterised [11]. Mutations in PA and PB1 have been found to contribute to higher polymerase activity and impact early viral replication, helping to expand the virus’ host range into humans [3, 19, 24]. Furthermore, a functional compatibility between PB1 and HA has been found to enhance viral fitness [25]. Viral fitness has been associated with the PB1 protein favouring more neutral and less reactive amino acids in certain positions.

## Discussion

Using genomic avian influenza data, we developed and applying a suite of MLAs to classify zoonotic transmission. We tested a number of models to determine that XGBoost is the optimal one. Applying XGBoost, we found that all 8 of its individual protein prediction models demonstrate strong predictive performance, particularly the HA and PB2 protein models. These findings are consistent with existing literature; other MLAs have also identified HA and PB2 as being the most important proteins in the prediction of zoonosis [26].

Furthermore, both of our combined prediction models demonstrate very strong predictive performance, with capability to differentiate between zoonotic and non-zoonotic avian influenza cases. In general, we see that the zoonotic cases which our combined model correctly predicts are also identified across numerous of our protein models. This is not exclusively the case, however, as some zoonotic cases are only identified by the combined model. This suggests that for zoonosis it is not just necessary for one protein to have a specific genetic feature. Instead, it is more likely that a multiple zoonotic genetic features across proteins are needed for zoonosis.

We have identified the secondary structure of HA, charge of PB2 and hydrophobicity of HA as some of the most important physiochemical features in our prediction model. We have suggested possible reasons why these physiochemical properties may be significant in prediction, but we believe that more research is needed to fully understand the role of these physiochemical properties.

We have shown that it is possible to classify avian influenza cases as zoonotic or non-zoonotic using XGBoost. When we use our combined models and our individual protein models in conjunction, we find that the most confirmed zoonotic cases have at least one zoonotic marker amongst all the models. We have shown that this majority can be as much as 98% when we set the minimum probability of a zoonotic marker as 0.01. This is promising for our prediction model as it shows that if we can measure the most significant genetic features for at least some of the proteins then we can use a combination of our models to accurately identify a case as zoonotic. If this can be done in a timely way, then we can identify avian influenza cases which have a high likelihood of being transmitted to humans and implement measures to prevent this.

Our results show how this could be done. We have identified a small number of non-zoonotic cases which have at least one zoonotic signature across the 10 models. Even in the most extreme cases when we set the minimum probability as 0.01, this is still less than 12% of all non-zoonotic cases. Figure 5 shows a visual example of a random selection 10 of these non-zoonotic cases. It shows the probability of being zoonotic that each of the 10 models assigns to these 10 non-zoonotic cases. For some of these non-zoonotic cases there are even multiple models with a notable probability of the case being zoonotic.

**Figure 5:**
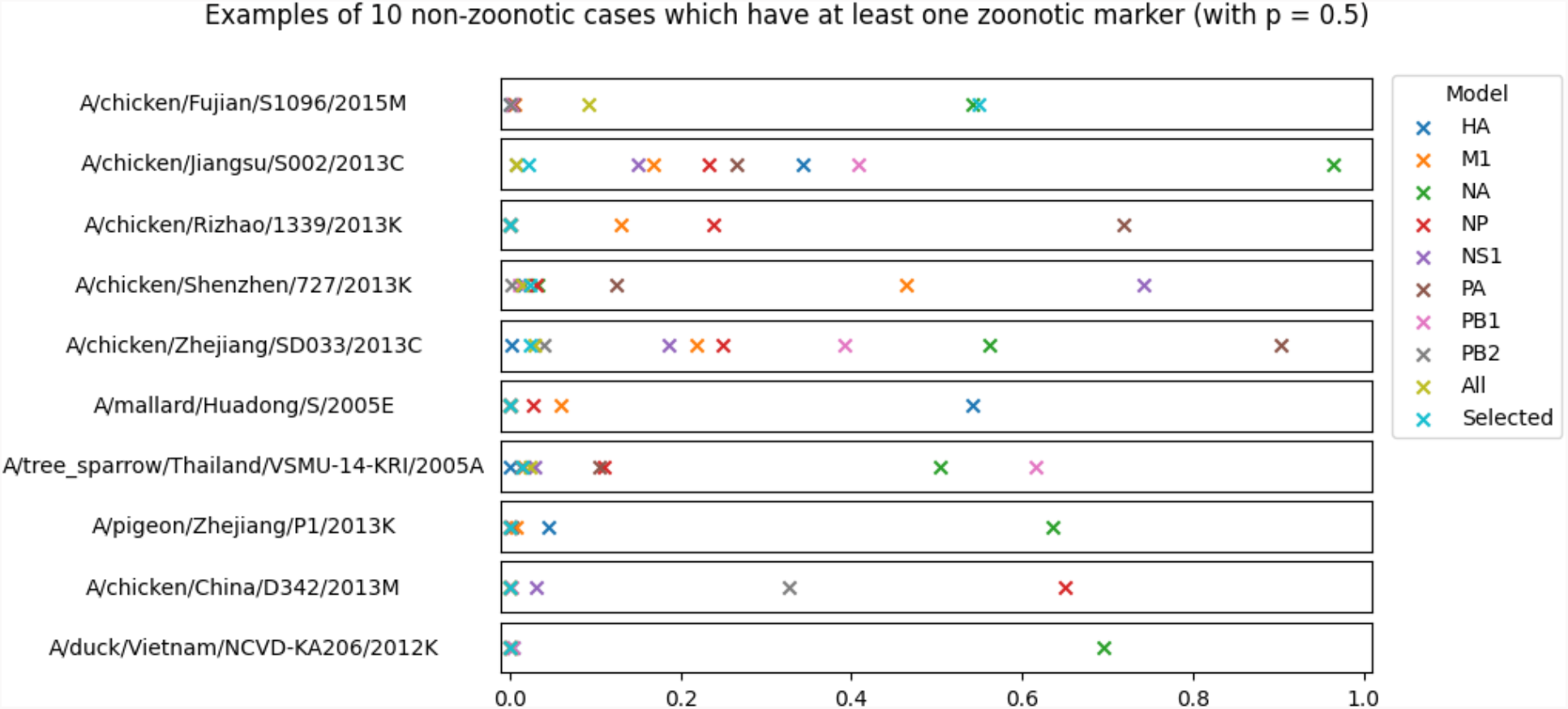
Examples of 10 randomly selected non-zoonotic cases found in avian hosts which have at least one zoonotic marker in one model (where we define a zoonotic marker as a model predicting zoonosis with more than 0.5 probability).

Let us consider the second case in Figure 5: A/chicken/Jiangsu/S002/2013C, an H7N9 avian influenza case found in a chicken host in Jiangsu in 2013. This avian case was identified as zoonotic by our NA model, with a high probability associated with this prediction. Within our dataset, we have some zoonotic cases that are the same subtype, from the same area and identified in the same year. Two examples of these are A/jiangsu/1/2013K A and A/jiangsu/1/2013K B. We have compared these cases in Figure 6. We observe that the NA model, amongst others, indicates a high probability that these human host cases are zoonotic. Given that all of these cases are from the same area in the same year, it is possible that the case found in the chicken (A/chicken/Jiangsu/S002/2013C) was part of the avian source which infected the humans.

**Figure 6:**
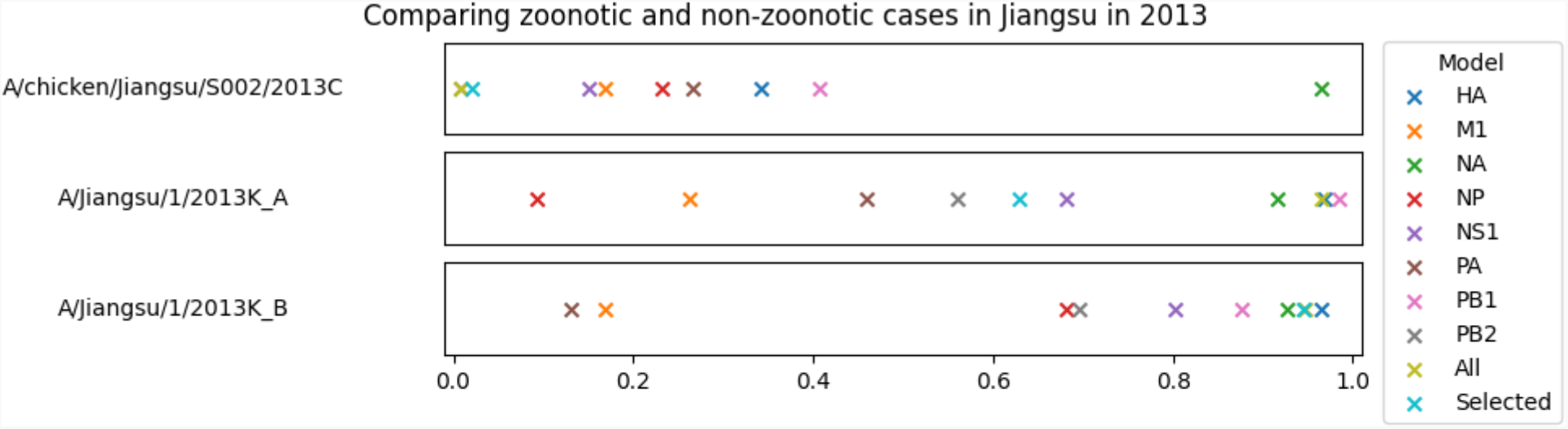
Comparing zoonotic markers between one non-zoonotic case of avian host (A/chicken/jiangsu/S002/2013C) and two zoonotic cases of human host (A/jiangsu/1/2013K A and A/jiangsu/1/2013K B) that are from the same subtype (H7N9), region (Jiangsu) and year (2013).

This is a clear example of exactly what we want our model to do: the model identifies the chicken (A/chicken/Jiangsu/S002/2013C) as a case that has risk of zoonotic potential through the NA protein model, and this same protein model is identifying zoonotic cases from the same area and time period correctly as zoonotic with similar probability. This supports an argument that the chicken case (A/chicken/Jiangsu/S002/2013C) is likely part of what caused the zoonotic avian influenza outbreak in this region in 2013. This shows that we can use our model to identify avian cases which could potentially lead to zoonotic cases, ideally before this spillover happens.

There are a number of limitations surrounding our model. Firstly, we have randomly split our data into a test and training set. In an ideal scenario we would want to split our data temporally, ensuring that all cases in the test data are occur at a later time than those in the training set. We attempted this temporal data split, but we were not achieve any results with reasonable performance as the data sets was not large enough for temporal split. Future work will look into combining the existing datasets with other data, and explore a temporal split further. Another limitation of our data is that only a few subtypes have a notable number of zoonotic cases. Relative to the total number of subtypes across the data set, only a small proportion of subtypes contain zoonotic cases. This could lead to our model performing significantly worse in a future avian influenza zoonotic spillover event if its subtype is one that our model did not have any zoonotic cases to train on. Hence, future works needs to use a larger dataset and potentially explore different methods to overcome this. Finally, our data is extremely geographically non-homogeneous. For example, there were no zoonotic cases (despite there being plenty on non-zoonotic cases) from the United States of America (USA) [27]. Considering that the USA has been a particular hotspot for avian influenza events in recent years, this is very likely to introduce bias into our model.

Overall, we anticipate that if we can measure genomic features in a timely way for new avian influenza outbreaks within avian host species, then we can measure their likelihood to make the zoonotic jump. Given limited resources, we suggest focussing on improved surveillance for infected birds that our prediction model identifies as sharing the closest genetic features to zoonotic cases. Before our model is rolled out as a predictive tool, we would like our model to have additional testing and external validation on more recent data as well as access to more spatially and temporally consistent data. This would help to overcome the limitations mentioned previously, reducing bias in our model and increasing its predictive ability. With this additional data, we would seek to investigate whether we could predict unseen variants/cases (through a temporally split test/training dataset) with similarly strong model performance. If possible, this would show that our model can predict new cases in real time, rather than modelling them retrospectively. This would open a platform for our approaches to be built in to future routine avian influenza surveillance.

Although in this study we have used the same dataset as in previous work by Brierley et al. [16], our work is an extension to this. Unlike that work, we used all the features in a single-model approach and we also outputted the SHAP values. Additionally, having undertaken a scoping review of existing models, we then compared a number of machine learning algorithms. Additionally, this work is an incremental building on the work completed by Eng et al. [28] and want to emphasise their influence on this work. Their approach of individual protein prediction models set the precedent for our combination of individual protein models and combined models. Their work also considered the probability assigned to each protein prediction model as a zoonotic risk indicator. We have similarly worked with probabilities to establish non-zoonotic cases of concern. Our work differs in two main areas: we use a wider variety of genetic features, and we use XGBoost as our prediction algorithm (as opposed to Random Forest). Eng et al. identified that the strength in any host tropism prediction system is found in its ability to predict false zoonotic cases. We fully agree with this and, similarly, suggest that our model’s strength lies in its ability to replicate this.

## 5. Conclusion

Our prediction models show that each protein can be a determinant of zoonosis, with clear differentiation between zoonotic and non-zoonotic cases and good model performance. We provide prediction models for 8 influenza proteins, and two combined models: one using all available genetic features and another using a significantly reduced list of genetic features. Both combined models have excellent performance, with the reduction in features only costing a small decrease in the performance of the model whilst significantly improving the model run time. We assert that our model can offer a fast, initial opinion on whether an AIV strain found within an avian host is likely to cross the species barrier in the near future.

## Data availability statement

Data for this study can be requested from the corresponding author, but restrictions apply to the availability of these data, which were used under license for the current study, and so are not publicly available.

## Acknowledgements

AF is supported by a PhD studentship from the UK Health Security Agency as part of the Oxford EPSRC Centre for Doctoral Training in Healthcare Data Science (EP/Y035321/1). JG and AA are supported by the EPSRC Centre for Doctoral Training in Healthcare Data Science (EP/Y035321/1). JPG’s work is supported by the UK Health Security Agency and the UK Department of Health and Social Care and the Oxford EPSRC Centre for Doctoral Training in Healthcare Data Science (EP/Y035321/1).

The funders had no role in the study design, data analysis, data interpretation, or writing of the report. The views expressed in this article are those of the authors and not necessarily those of the UK Health Security Agency, the UK Department of Health and Social Care or the EPSRC Centre for Doctoral Training in Healthcare Data Science.

## Conflict of Interest

None

## Authors contribution

Formal analysis: AF, AA and JG; Validation: AF; Writing – original draft: AF and JPG; Writing – review & editing: LM, AA, JG, LC; Data curation: AF and LB; Methodology: AF, AA, JG, LM and JPG; Visualization: AF; Investigation: AF, LB and JPG; Conceptualization: JPG; Software: AF, AA and JG; Supervision: JPG.; Project administration: AF and JPG.

## Code availability

The numerical code used in this analysis can be requested from the corresponding author. Restrictions apply to the availability of the model combining code, which was under collaborative license for the current study and hence is not publicly available.

## Notes

### Competing Interest Statement

The authors have declared no competing interest.

