## Appendix for "Predicting the Risk of Avian Influenza Zoonosis using Viral Genome Sequencing Data"

### A. Appendix

*A.1. Literature Review*

We used the following search criteria with an unrestricted date range in PubMed: ((influenza[Title/Abstract]) OR (avian influenza[Title/Abstract])) AND ((((zoonotic[Title/Abstract])) OR (human[Title/Abstract])) AND (transmission[Title/Abstract])) OR (host tropism[Title/Abstract]) OR (zoonosis[Title/Abstract]))) AND ((machine learning[Title/Abstract]) OR (deep learning[Title/Abstract]) OR (unsupervised learning[Title/Abstract]) OR (supervised learning[Title/Abstract])). This yielded 53 results. 38 of these were removed after reading their abstracts due to irrelevance to our work. The remaining 15 papers could be split into different categories.

Eleven papers consider Machine Learning (ML) algorithms, and another five look at Deep Learning (DL) algorithms (one paper includes both ML and DL methods). We identified an additional 9 papers relevant to our work through the citations of the previous 15 papers. Seven of these consider ML algorithms and three consider DL algorithms (one paper includes both ML and DL methods). We present a summary of the algorithms used and predictive ability achieved by these papers, beginning with DL and then moving to ML. A summary of the key elements of all 23 reviewed papers can be found in Table A.5.

Deep learning of influenza protein genomic sequences is utilised across papers to make viral host predictions. Mock et al. [29] used deep neural networks to assign one of 36 different host species to genome sequences of influenza. Similarly, Qiang and Kou [30] predicted interspecies transmission of AIV using a BP neural network based on wavelet packet decomposition that transformed the genome sequences into energy features that were used as the model input. Some work has used convolutional neural networks (CNN). Borkenhagen and Runstadler [31] developed a CNN to predict the $\alpha$2,6-linked SA preference of IAV given the amino acid sequence of the HA protein. They found that the highest SHAP values were localised to the globular head of HA rather than the stalk region, in line with biological understanding. They removed all samples of one strain and still had good, albeit decreased, performance showing that the model could be used in a future prediction setting with unknown strains.

Chrysostomou et al. [32] used the amino acid sequence of the HA protein to predict whether the host species is one of the 3 major hosts: avian, swine or human. This tri-category prediction method was popular across papers. They achieved unexpectedly high performance, as measured by the total accuracy, MCC and F1 score. In contrast, Hatibi et al. [33] used two different deep learning models (BiDirectional LSTM and Transformers). They used the amino acid sequence and mRNA sequence on tri-category prediction with multiple AIV proteins and found that the mRNA sequence led to better performance than the amino acid sequences. Their results suggest that codon sequences contain important information about viral hosts that is lost in translation to amino acids. This was further verified by the specific codons contributing to changes in the likelihood of zoonotic transmission. They hypothesised that sequences which are difficult to classify are the best candidates for zoonotic transition, as they contain biomarkers indicative of zoonosis. Their work offers evidence in support of this hypothesis.

Other papers have compared both DL and more traditional ML and statistical methods to predict host tropism. Scarafoni et al. [34] used influenza amino acid sequences of multiple proteins from avian and human samples to identify potential zoonotic candidates. They found a tailored CNN to be the best performer due to the end-to-end learning it completes on the sequences. Their analysis did not factor the phylogenetic similarities between some strains, assuming that they were all independent, meaning that their model may not have good predictive capabilities for future unknown strains. Xu and Wojtczak [35] also used CNN to determine host tropism, although they only looked at the HA protein. They considered alignment-based (i.e. genetic sequence) and alignment-free (frequency measurements) approaches to train their model. They found their transformer neural network to outperform other models in most scenarios. Other studies that compare alignment-based and alignment-free approaches have been done by Li and Sun [36]. They look at a support vector machine (SVM) approach, this time using mononucleotide frequency and dinucleotide bias.

| Ref. | ML/DL | Algorithms used | Features used | Proteins included | Classification goal |
|---|---|---|---|---|---|
| [29] | DL | Deep NN | Viral genome fragments (100–400bp) | Viral genome fragments | Host species |
| [30] | DL | BPNN, Wavelet Package Decomposition | (Unspecified) | M1, NP, NS1, PA, PB1, PB2 | Zoonosis within H or A host |
| [31] | DL | CNN | AA sequence (one-hot or embeddings) | HA | SA binding preference |
| [32] | DL | CNN | AA sequence | HA | H, A or S host |
| [33] | DL | BiLSTM, Trans. | mRNA sequence, AA sequence | (Unspecified proteins) | H, A or S host |
| [34] | ML, DL | CNN | AA sequence | HA, M, M1, NEP, PA, PB, PB1-F2 | Zoonosis within H or A host |
| [35] | ML, DL | RF, RUSboost, SVM, XGB, MLP, Transformer, CNN | Sequence-derived features (e.g., position-specific scoring matrix, k-mer, embeddings) | HA | (Classification goal unspecified) |
| [36] | ML | SVM | (Di)nucleotide composition, frequency, bias | Whole sequence, not protein specific | Host species |
| [37] | ML | DT, NB, RF, SVM | AA sequence, AAFactors | (Unspecified proteins) | Interspecies transmission |
| [38] | ML | SVM, RF, KNN, NB | Sequence-derived features | HA, M1, M2, NA, NEP, NP, NS1, PA, PB1, PB1-F2, PB2 | Interspecies transmission |
| [26] | ML | SVM, KNN, NB | Sequence-derived features | HA, M1, M2, NA, NEP, NP, NS1, PA, PB1, PB1-F2, PB2 | H or A host |
| [39] | ML | CBA, Ripper, DT | AA sequence features | HA, M1, M2, NA, NP, NS1, NS2, PA, PA-X, PB-1, PB1-F2, PB2 | H, A or S host |
| [40] | ML | RF | AA sequence features | (Unspecified proteins) | (Classification goal unspecified) |
| [41] | ML | HMM, DT, Associative Classification | AA sequence features | HA | H, A or S host |
| [42] | ML | (Unspecified) | Clinical and virological features | HA | (Classification goal unspecified) |
| [43] | ML | LR, KNN, RF, GNB, SVM, MLP | (K)nucleotide frequency | HA, NA, MP, NP, NS, PA, PB1, PB2 | (Classification goal unspecified) |
| [44] | ML | GBRT, MLP, RF, SVC | (Di)nucleotides | HA, NA, NP, PA, PB1, PB2 | H, A or S host |
| [45] | ML | SVM, RF, KNN, DT, SGB, GNB | AA composition, DPC | HA, M1, M2, NA, NP, NS1, NS2, PA, PA-N155, PA-N182, PA-X, PB1, PB1-F2, PB1-N40, PB2 | Zoonosis within H or A host |
| [46] | ML | LR, kNN, GBM, RF | (K)nucleotide, codon pair score/bias, AA composition/properties | HA | (Classification goal unspecified) |
| [47] | ML | (Unspecified) | Physicochemical properties | HA | (Classification goal unspecified) |
| [28] | ML | | CTD features | HA, M1, M2, NA, NP, NS1, NS2, PA, PB1, PB1-F2 | H or A protein signatures |
| [48] | ML | Hierarchical clustering | CTD features | HA, M1, M2, NA, NP, NS1, NS2, PA, PB1, PB1-F2 | Binary host prediction model |
| [49] | ML | Computational model | CTD features | HA, M1, M2, NA, NP, NS1, NS2, PA, PB1, PB1-F2 | Probability-based host prediction model |

Table A.5: Summary table of all papers included in the literature review.

Li and Sun [36] focussed on predicting one of six host species. This is similar to the host species model developed by Mock et al. [29]. They did not find any of their algorithms to perform particularly well, with prediction accuracies below 0.6.

Several other papers have used alignment-based and alignment free methods in the form of signature amino acid positions within sequences combined with amino acid factor scores (AAFactors) to predict inter-species transmission of AIV [37, 38]. AAFactors concentrate many physiochemical and biochemical properties of amino acids into five factor scores: polarity, secondary structure, molecular volume, codon diversity and electrostatic charge. Wang et al. [37] used various ML algorithms: Decision Tree (DT), Naive Bayes (NB), Random Forest (RF) and SVM to integrate these AAFactors with signature amino acid positions previously identified. Using similar ML algorithms (NB, RF, SVM and k-Nearest Neighbours (kNN)) on multiple proteins, Qiang et al. [38] found that the HA protein had the largest number of signature amino acid positions, followed by NA. These positions were in important functional regions of the receptor binding site. They further found that molecular volume was the most important AAFactor. SVM was the overall best performer out of all their models.

Following similar methods, Qiang and Kou [26] use the AAFactors and amino acid positions to classify AIV cases into that of human-origin and avian-origin. They use NB, SVM and kNN on multiple proteins and find HA and PB2 to have the most important roles in prediction.

A large number of the ML algorithms have focussed on sequence alignment-based methods alone. Kargarfard et al. [39] used classification based on association (CBA) algorithm, Ripper algorithm, and DT to identify amino acid positions within multiple proteins that determine the classification of AIV cases into one of the three major hosts. They found multiple genes (HA, NS1, NA and PB1) to have specific amino acid positions with an importance of more than 10%. ElHefnawi and Sherif [41] also classified AIV into the three major hosts, but used only the HA protein. They considered Hidden Markov Models (HMMs), DTs and CBA. They identified amino acid residues within the HA protein that were specific to either human, swine or avian influenza viruses. Aguas and Ferguson [40] similarly used specific amino acid residues and substitutions, this time using RF to classify AIVs into their host species reservoir. They found certain substitutions were only determinant when found in conjunction with certain substitutions at other sites.

In contrast, other papers have focussed solely on alignment-free methods. An outlier amongst all other papers considered is [42]. They used ML techniques on molecular determinants, clinical parameters and infectious titer metrics of the HA gene segment in transmission events in ferrets.

Sun et al. [43] used weight-based Logistic Regression (LR) kNN, RF, Gaussian Naive Bayes (GNB), SVM and Multi-Layer Perceptron (MLP) to find specific nucleotide sequences for multiple proteins which are predictors of zoonotic transmission for the H7N9 AIV strain. They were able to recognise the human H7N9 strain genome with 100% accuracy using the weight-based LR. Mononucleotide and dinucleotide composition in multiple proteins were also used to inform an ML model by Li et al. [44]. They used Principal Component Analysis (PCA), Hierarchical Clustering Analysis (HCA) and Support Vector Classification (SVC) to find linear separability of optimised (di)nucleotides. They then used gradient boosted regression trees (GBRT), MLP, RF and SVC to classify AIV cases into the three major hosts. They found that between 9 and 13 (di)nucleotides is enough to predict human adaptation for each of the protein segments.

Other work has looked at the amino acid compositon rather than nucleotide composition. Roy et al. [45] have used ML algorithms (SVM, RF, kNN, DT, XGBoost (XGB) and GNB) to find specific amino acid compositions and dipeptide compositions (DPCs) that are indicative of zoonosis for multiple AIV proteins. They found RF to be the best performing model for this task. Similarly, Alberts et al. [46] predict the most likely host species using amino acid frequency, (di)nucleotide frequency, 4-mer, 5-mer, codon frequency, codon pair score, codon pair bias and amino acid properties. Unlike Roy et al. [45], they only consider these features for the HA protein. The amino acid properties included hydrophobicity, polarity and net charge. They utilised the per Gini score to determine the top 10% of features after using RF on the training set. They found that most of the selected features were based of the nucleic acid sequence rather than the amino acid sequences/properties.

On the other hand, the importance of the physiochemical properties of amino acids has been focussed on elsewhere. Yin et al. [47] have used RF on the AIV HA protein to predict host tropism, based on physiochemical properties that influence the selection of binding to different hosts. They considered seven different physiochemical properties of amino acids. They used feature aggregation and associative rules to select the top 20 features. They found that the secondary structure and normalised van der Waals volume were most important in determining host tropism.

The final model we discuss is a combination of three papers [28, 48, 49]. Amongst all the relevant literature, we have concluded that these studies provide the most complete end-to-end process of how ML algorithms can be used to create an accurate and rapid prediction model that can aide early detection of AIV cases at risk of becoming zoonotic. Initially, Eng et al. [28] used traditional ML algorithms to identify CTD features of different influenza proteins that are indicative of zoonosis. They had data from human and avian IAV samples in approximately equal numbers for 12 different proteins. Feature selection was done using RF and per Gini score, similar to Roy et al. [45]. They took the top 15 features for each protein and then included these in their combined prediction model, which was constructed to predict influenza virus host (avian or human) using composition-transition-distribution (CTD) features of each protein. For the HA gene, they found that charge, normalised van der Waals and polarizability were the most significant features. They identified that the strength of any host tropism prediction system lies not in perfect prediction accuracy, but actually in its mistakes in classifying avian cases as human.

Eng et al. [48] continued this work by performing hierarchical clustering on their protein signatures once they had identified suspected and confirmed zoonotic cases. They set a condition of an avian strain needing at least one 'human' protein signature to be a suspected zoonotic case, which was less than 4% of all avian strains. The idea behind this was that an avian strain isn't capable of crossing the host species barrier without at least one 'human' protein signature. The clustering showed that strains isolated from the same outbreak shared similar human protein signatures. They hypothesised that as avian strains have more human features, they can eventually transmit zoonotically. In the confirmed zoonotic strains, five of the proteins (M2, NA, NS1, PA, PB1-F2) had more than 50% human predictions. Surprisingly the well-known protein determinants of host adaptation, HA and PB2, were not among the top proteins with the most human predictions. At least 1/3 of all confirmed zoonotic strains had between 5-10 human proteins (out of 12). This shows that the AIV does not need all human protein signatures to have zoonotic capability.

The final paper following on with this work aimed to build a computational model to predict zoonotic strains. Eng et al. [49] decided to turn their problem into a three-class classification problem by adding zoonotic strains as a distinct category from avian and human strains. Instead of using binary predictions for each protein signature, they had avian and human probability distributions. The probabilistic nature of RF allowed their model to have an avian, human and zoonotic probability estimate. They found that some zoonotic strains carry all avian protein signatures when taken at binary level, yet were predicted accurately as zoonotic strains by their combined prediction model. This suggests that zoonotic strains don't need to acquire human protein signatures to transmit zoonotically, contradicting their previous theory.

The power of this work lies in its potential to recognise zoonotic strains regardless of subtype or specific amino acid residues. This can allow for rapid prediction of potential zoonotic AIV candidates which can then be further analysed for their host-associated genomic markers. Furthermore, they identified that if they could include geographical data in their future work, it would bring more significant advancements in predicting potential future avian influenza outbreaks, potentially outperforming current sequence prediction capabilities.

*A.2. Genomic Feature Definitions*

*A.2.1. Nucleic Acid Composition*

Nucleic Acid Composition (NAC) of length 2 for a nucleic acid sequence $x_1, x_2$ where $x_i$ is one of the 4 nucleic acids A (adenine), G (guanine), C (cytosine), T (thyamine) is defined as:

$$\text{NAC}(x_1, x_2) = \frac{n_{x_1,x_2}}{N},$$

where $n_{x_1,x_2}$ is the number of appearances of the nucleic acid sequence $x_1, x_2$ and $N$ is the total number of nucleic acid sequences of length 2. There are 64 nucleic acid composition features per protein.

*A.2.2. Dipeptide Composition*

Dipeptide Composition (DPC) is defined as:

$$\text{DPC}(r, s) = \frac{N_{r,s}}{N-1},$$

where $r, s \in \{A, C, D, ..., Y\}$ and $N_{r,s}$ is the number of dipeptides represented by amino acids type $r$ and $s$ and $N$ is the length of the peptide sequence. There are 400 ($20 \times 20$) DPC descriptors within the dipeptide feature group.

*A.2.3. Composition-Transition-Distribution*

The Composition-Transition-Distribution (CTD) features measure AA distribution patterns of specific structural and physiochemical properties in a protein. There are 13 types of these properties: hydrophobicity (7 different types), normalized Van der Waals Volume, polarity, polarisability, charge, secondary structures and solvent accessibility. All AAs are divided into 3 groups (as shown in Table A.6, which vary depending on which physiochemical property you are interested in.

| **Physiochemical Property** | $G_1$ | $G_2$ | $G_3$ |
|---|---|---|---|
| Hydrophobicity_PRAM900101 | Polar: RKEDQN | Neutral: GASTPHY | Hydrophobicity: CLVIMFW |
| Hydrophobicity_ARGP820101 | Polar: QSTNGDE | Neutral: RAHCKMV | Hydrophobicity: LYPFIW |
| Hydrophobicity_ZIMJ680101 | Polar: QNGSWTDERA | Neutral: HMCKV | Hydrophobicity: LPFYI |
| Hydrophobicity_PONP930101 | Polar: KPDESNQT | Neutral: GRHA | Hydrophobicity: YMFWLCVI |
| Hydrophobicity_CASG920101 | Polar: KDEQPSRNTG | Neutral: AHYMLV | Hydrophobicity: FIWC |
| Hydrophobicity_ENGD860101 | Polar: RDKENQHYP | Neutral: SGTAW | Hydrophobicity: CVLIMF |
| Hydrophobicity_FASG890101 | Polar: KERSQD | Neutral: NTPG | Hydrophobicity: AYHWVMFLIC |
| Normalized Van der Waals Volume | Volume range: 0-2.78 GASTPD | Volume range: 2.95-94.0 NVEQIL | Volume range: 4.03-8.08 MHKFRYW |
| Polarity | Polarity value: 4.9-6.2 LIFWCMVY | Polarity value: 8.0-9.2 PATGS | Polarity value: 10.4-13.0 HQRKNED |
| Polarizability | Polarizability value: 0-0.108 GASDT | Polarizability value: 0.128-0.186 GPNVEQIL | Polarizability value: 0.219-0.409 KMHFRYW |
| Charge | Positive: KR | Neutral: ANCQGHILMFPSTWYV | Negative: DE |
| Secondary Structure | Helix: EALMQKRH | Strand: VIYCWFT | Coil: GNPSD |
| Solvent Accessibility | Buried: ALFCGIVW | Exposed: PKQEND | Intermediate: MPSTHY |

Table A.6: Division of amino acids into 3 groups based on their physiochemical properties.

*Composition*

The CTD Composition (CTDC) for a specific physiochemical property is defined as:

$$\text{CTDC}(r) = \frac{N(r)}{N}, \quad r \in \{G_1, G_2, G_3\},$$

where $N(r)$ is the number of amino acids of type r in the sequence, and $N$ is the length of the sequence.

*Transition*

The CTD Transition (CTDT) for a specific physiochemical property is defined as:

$$\text{CTDT}(r) = \frac{N(r,s) + N(s,r)}{N-1},$$

$r, s \in \{(G_1, G_2), (G_1, G_3), (G_2, G_3)\}$ where $N(r,s)$ is the number of dipeptides encoded as $rs$ and $sr$ respectively in the peptide sequence, and $N$ is the length of the sequence.

*Distribution*

The CTD Distribution (CTDD) consists of five values for a specific physiochemical property and group ($G_1, G_2$ or $G_3$). Each value corresponds to a different residue (0, 25, 50, 75, 100%). The first value corresponds to the fraction of the entire peptide sequence included up to the first residue of a given group. The following values correspond to the proportion of the total length of the sequence where 25, 50, 75 and 100% of occurrences are contained.

*A.2.4. Conjoint Triad*

To calculate the Conjoint Triad (CTriad) we first categorise each amino acid into 7 different categories (as shown in Table A.7). We then consider all groups of three consecutive amino acids as a single unit that is made up of 3 elements, each belonging to one of the 7 categories. Thus, there are 343 ($7 \times 7 \times 7$) CTriads. The CTriad is defined as:

$$\text{CTriad}(g_i, g_j, g_k) = \frac{N(g_i, g_j, g_k)}{D_{ijk}},$$

where $N(g_i, g_j, g_k)$ is a the number of amino acids triads of the form $(g_i, g_j, g_k)$ for $i, j, k \in \{1, 2, ..., 7\}$ and $D_{ijk}$ is a normalising constant. We define $D_{ijk}$ as:

$$D_{ijk} = \frac{N(g_i, g_j, g_k) - \min\{N(g_l, g_m, g_n)}{\max\{N(g_l, g_m, g_n)},$$

$\forall l, m, n \in \{1, 2, ..., 7\}$ and $l \neq i, m \neq j, n \neq k$.

| Group | Amino Acids |
|---|---|
| $g_1$ | A,G,V |
| $g_2$ | I,L,F,P |
| $g_3$ | Y,M,T,S |
| $g_4$ | H,N,Q,W |
| $g_5$ | R,K |
| $g_6$ | D,E |
| $g_7$ | C |

Table A.7: Division of amino acids into 7 groups.

*A.2.5. Pseudo-Amino Acid Composition*

Given a protein sequence of length $L$, the Pseudo-Amino Acid Composition (PAAC) is a vector $\mathbf{P}$ given by:

$$\mathbf{P} = [p_1, p_2, \ldots, p_{20}, p_{21}, \ldots, p_{20+\lambda}],$$

where:

- $p_1, p_2, \ldots, p_{20}$ are the normalised occurrence frequencies of the 20 standard amino acids (A, R, N, D, C, E, Q, G, H, I, L, K, M, F, P, S, T, W, Y, V);
- $p_{21}, \ldots, p_{20+\lambda}$ are the sequence-order correlation factors (pseudo components);
- $\lambda$ is a user-defined parameter indicating the highest rank of sequence correlation considered ($\lambda \leq L-1$).

We first calculate the normalised amino acid composition components:

$$p_u = \frac{f_u}{1 + w \sum_{k=1}^{\lambda} \theta_k}, \quad u = 1, 2, \ldots, 20,$$

where $f_u = n_u/L$, $n_u$ is the count of the $u^{\text{th}}$ amino acid in the sequence, and $w$ is a weight factor balancing the effect of sequence order information relative to amino acid composition.

We need calculate the sequence order correlation factors $\theta_k$:

$$\theta_k = \frac{1}{L-k} \sum_{i=1}^{L-k} \Theta(R_i, R_{i+k}), \quad k = 1, 2, \ldots, \lambda,$$

where $R_i$ is the amino acid residue at position $i$.

We then define the correlation function $\Theta$:

$$\Theta(R_i, R_j) = \frac{1}{m} \sum_{t=1}^{m} \left[ H_t(R_i) - H_t(R_j) \right]^2,$$

where $m$ is the number of selected physicochemical properties and $H_t(R_i)$ is the normalized value of the $t$-th physicochemical property for residue $R_i$.

Finally, we calculate the pseudo components $p_{20+k}$:

$$p_{20+k} = \frac{w\theta_k}{1 + w \sum_{k=1}^{\lambda} \theta_k}, \quad k = 1, 2, \ldots, \lambda.$$

We choose $w = 0.05$, $\lambda = 3$ and $m = 3$, to account for the hydrophobicity value, hydrophilicity value and side chain mass.

*A.3. Machine Learning Performance Measure Definitions*

In our binary classification problem we define:

- a correctly classified zoonotic case as a true positive (TP);
- an incorrectly classified zoonotic case as a false positive (FP);
- a correctly classified non-zoonotic case as a true negative (TN);
- an incorrectly classified non-zoonotic case as a false negative (FN).

We use a number of performance measures, with the following definitions:

1. Accuracy is the total proportion of predictions which are correct.

$$\text{Accuracy} = \frac{TP + TN}{TP + TN + FP + FN}$$

2. Precision is the proportion of predicted positives that are actually positive.

$$\text{Precision} = \frac{TP}{TP + FP}$$

3. Recall is the proportion of actual positives that are correctly identified.

$$\text{Recall} = \frac{TP}{TP + FN}$$

4. F1 score is the harmonic mean of precision and recall.

$$\text{Precision} = \frac{2TP}{2TP + FP + FN}$$

5. Specificity is the proportion of actual negatives that are correctly identified.

$$\text{Specificity} = \frac{TN}{TN + FP}$$

6. Matthew's Correlation Coefficient (MCC) is a correlation coefficient between observed and predicted classifications that ranges from -1 (total disagreement) to +1 (perfect prediction).

$$\text{MCC} = \frac{TP \cdot TN - FP \cdot FN}{\sqrt{(TP + FP)(TP + FN)(TN + FP)(TN + FN)}}$$

7. Area Under the ROC Curve (AUC) measures the area under the receiving operating characteristic (ROC) curve which plots recall against the false positive rate ($FPR = 1 -$ specificity) at various thresholds, with values ranging from 0.5 (random) to 1.0 (perfect).

$$\text{AUC} = \int_0^1 \text{recall} \cdot (FPR)\ d(FPR)$$

8. Average Precision is the average value of precision across different recall values.

$$\text{Average Precision} = \int_0^1 \text{precision}(\text{recall})\ d(\text{recall})$$
